# Paraventricular thalamus gates hippocampal coding of salient experiences

**DOI:** 10.64898/2026.05.29.728856

**Authors:** Mark M. Gergues, Gregory I. Telian, Kelsey R. Logas, Mona Li, Arsine Kolanjian, Jeremy S. Biane, Shazreh Hassan, Jasmine Tai, Victoria S. Turner, Mazen A. Kheirbek

## Abstract

Adaptive behavior requires neural circuits that encode salient stimuli to guide approach and avoidance. The hippocampal ventral subiculum (vSub) encodes information related to avoidance and reward seeking, yet how upstream inputs shape its representations of salient appetitive and aversive events remains unclear. Here, we show that a projection from the anterior paraventricular thalamus (PVT) to vSub carries an aversive-state signal that biases vSub representations toward threat-related information. In anxiety-provoking environments, anterior PVT to vSub activity is associated with avoidance behavior and enhanced vSub discrimination of threat and safety. During associative learning, this circuit preferentially promotes vSub responses to threat-predictive cues and aversive outcomes while constraining representations of rewarding stimuli. These findings refine canonical limbic circuit models by identifying a direct thalamic influence over hippocampal coding of rewarding versus aversive stimuli and the approach–avoidance behaviors they guide.

## Introduction

During exploration, animals must balance reward seeking with threat avoidance, guided by representations of emotionally salient experiences and contexts. The ventral hippocampus (vHPC) is a key node in this process, responding to emotionally salient events and shaping fear, anxiety, and reward-related behaviors through its interactions with downstream targets such as the medial prefrontal cortex, lateral hypothalamus, nucleus accumbens, and amygdala^1–9^. vHPC activity is modulated during exploration of rewarding and threatening environments, and perturbing vHPC alters anxiety-like behavior and reward seeking via these pathways^10–12^. Recent work further shows that vHPC neurons discriminate among stimuli, both innately salient and those that acquire salience through learning^1,2,7^. A central open question is how information about threat- and reward-associated stimuli, together with the internal and behavioral states they evoke, is routed into vHPC, and which upstream inputs tune its coding of motivational experiences. Previous work has shown that upstream amygdala neurons projecting to the vHPC encode cues that predict reward and aversion^13,14^, and manipulation of those amygdala axons in the vHPC can drive avoidance behavior^15^. However, whether this directly contributes to how vHPC encodes rewarding and salient events, and if it is the sole upstream contributor or how other upstream areas could contribute to vHPC coding, remains to be elucidated.

Canonical limbic models such as the Papez circuit^16,17^ have focused on indirect polysynaptic pathways between the hippocampus and thalamus and have not incorporated direct thalamic input to vHPC output circuits, obscuring a route by which the thalamus could shape affective salience representations and bias approach and avoidance decisions. Recent anatomical studies have identified a specific projection from anterior PVT (PVTa) to vHPC^18–22^, with especially strong input to vHPC neurons that represent aversive states and project to the lateral hypothalamus. More broadly, the PVT has recently attracted greater interest^23–25^ for its role as a limbic hub that integrates arousal, reward, fear, stress, and interoceptive signals through its cortical and subcortical connections^26,22,27,20,28,29,25,30–45^. However, its functional role in tuning vHPC coding and behavior is unknown. Here, we tested whether a PVTa projection to the subicular subregion of vHPC (vSub), which receives the densest innervation from PVTa, relays information about appetitive and aversive stimuli and internal state. To do so, we use high-throughput single-neuron tracing, optogenetic and chemogenetic perturbations, high-density electrophysiology, freely moving miniature microendoscopy, and two-photon calcium imaging to define the organization of this pathway and its impact on vSub population activity during associative learning and exposure to rewarding, safe, and aversive outcomes.

## Results

### Single neuron organization of anterior PVT projections to ventral hippocampus

To map the projection patterns of individual PVT neurons, we used high-throughput single-neuron tracing with multiplexed analysis of projections by sequencing (MAPseq)^18,46–48^. Posterior PVT neurons exhibit extensive collateralization^33,39,49,50^, but whether anterior PVT neurons show similar connectivity when assessing more than 3 downstream regions and how vSub-projecting neurons are organized was unclear. We injected a library of RNA barcodes into the PVTa, so that each neuron incorporated a unique barcode that was transported down the axon. We dissected the injection site and seven anatomically defined target regions (vSub, mPFC, NAc, bed nucleus of the stria terminalis (BNST), central amygdala (CeA), zona incerta (ZI), and BLA) and sequenced barcodes in each area to infer single-neuron projection patterns (Fig. 1A-B), using the relative abundance of each barcode in each region as a proxy for projection strength. We profiled projections from over 1,800 PVTa neurons and found that most (60.2%) projected to one of the assayed targets, with 30.2 % of these projecting exclusively to vSub (Fig. 1C, Supplemental Table 1). In addition, a substantial fraction of the total population of neurons collateralized to multiple targets: 31.2% projected to two regions, 7.2 percent to three, 1.1% to four, and 0.29% to five or more (Fig. 1C-D). In fact, when looking at the conditional probability of PVT neurons projecting to each region, we find hot spots of high likelihood of collateralization among NAc- and BNST-projecting neurons (as previously seen with retrograde studies^33,39,50^) and vSub (Fig. 1E).

**Figure 1:**
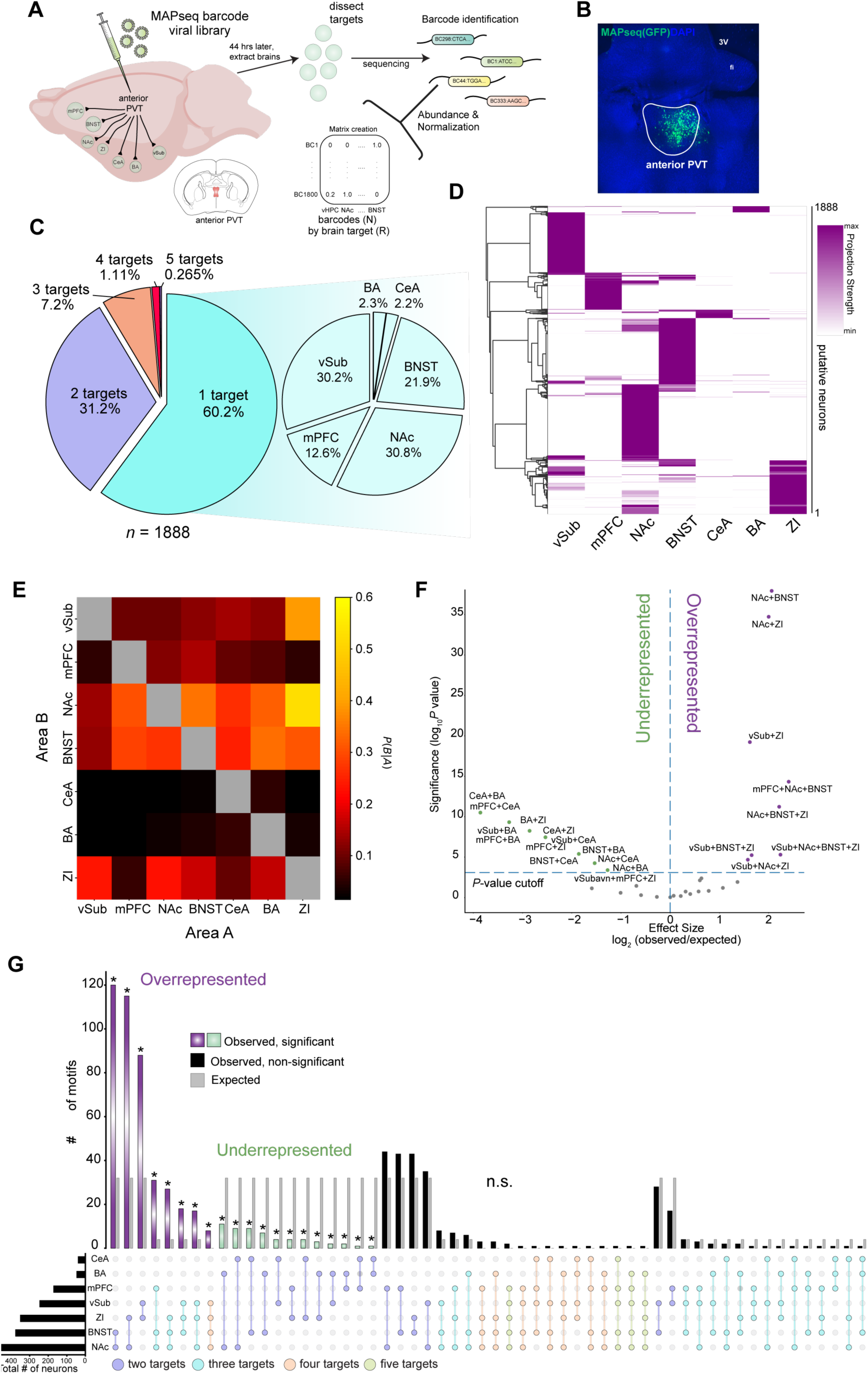
Single cell logic of PVTa projections. **A.** Experimental design of MAPseq barcode viral library injections into the PVT, after ∼44-46 hours following surgery, microdissections of the PVT, mPFC, NAc, BNST, CeA, BLA, ZI, and vHPC were collected from *n* = 9 mice and sent for RNA sequencing to identify and analyze barcodes. **B.** Example image of barcode-expressing GFP-tagged neurons at the injection site in PVT. **C.** Proportion of projection type and the breakdown of single-projectors by area. **D.** Heatmap of the profiled 1888 neurons in PVT across the 7 sampled areas, with intensity in purple depicting normalized relative projection strength for each barcode. **E.** Heat map of conditional probability for two regions, indicating the proportion of cells projecting to area B within the subset of cells that project to area A. **F.** Effect size and significance cut-offs for all motifs in dashed lines and grey circles. Over-represented motifs are colored in purple and under-represented motifs are green. **G.** Upset plot showing the number of motifs in observed data compared against the null distribution. Gray bars represent expected counts, with the mean and standard deviation of the null binomial models *P* values were calculated from two-sided binomial tests, adjusted for multiple comparisons using the Bonferroni method; * *P*_adj_ < 0.05.

To examine how bifurcating PVTa neurons are organized, we quantified projection motifs, defined as unique combinations of downstream regions innervated by individual neurons, and compared their frequencies to a null model in which projections to each target occurred independently (Fig. 1F-G). This analysis revealed a small set of over-represented motifs, most prominently neurons co-projecting to nucleus accumbens (NAc) and bed nucleus of the stria terminalis (BNST), and several under-represented motifs, including neurons projecting to both basal amygdala (BA) and central amygdala (CeA). Within the vSub-projecting population, most PVTa neurons projected exclusively to vSub (58.1%), with smaller fractions collateralizing to combinations including ZI, BNST, and NAc, areas without dense direct input back to vHPC^18^. These patterns were corroborated using a synaptophysin-based anterograde tracing approach (Supplementary Fig. 1). Together, these data indicate that the PVTa sends a dense projection to vSub composed largely of vSub-only neurons.

### PVTa population activity tracks anxiety-related exploration

We next asked how PVTa neurons are engaged during exploration of an anxiogenic environment. We performed high-density single-unit recordings with chronically implanted Neuropixels 2.0 probes targeting PVTa while mice explored an elevated plus maze (EPM; Fig. 2A). Across the population, a larger fraction of neurons showed higher firing rates in the open arms than in the closed arms, indicating an enrichment of units selective for the anxiogenic open arms (Fig. 2B-D). Examining single-neuron activity of open selective neurons aligned to trajectories from the closed to the open arms (Fig. 2E), we identified a subset of PVTa neurons that increased firing at entry into the open arms. A further subset showed peak firing when mice engaged in head-dip risk assessment at the edge of the open arms, the most aversive portion of the maze (Fig. 2G-F). At the population level, arm identity could be decoded from neural activity with accuracy significantly above shuffled controls (Fig. 2H). These data indicate that PVTa neurons encode aversive spaces and are highly responsive during highly anxiogenic risk-assessment behaviors during exploration of the EPM, providing single-cell evidence that PVTa carries detailed information about aversive states.

**Figure 2.**
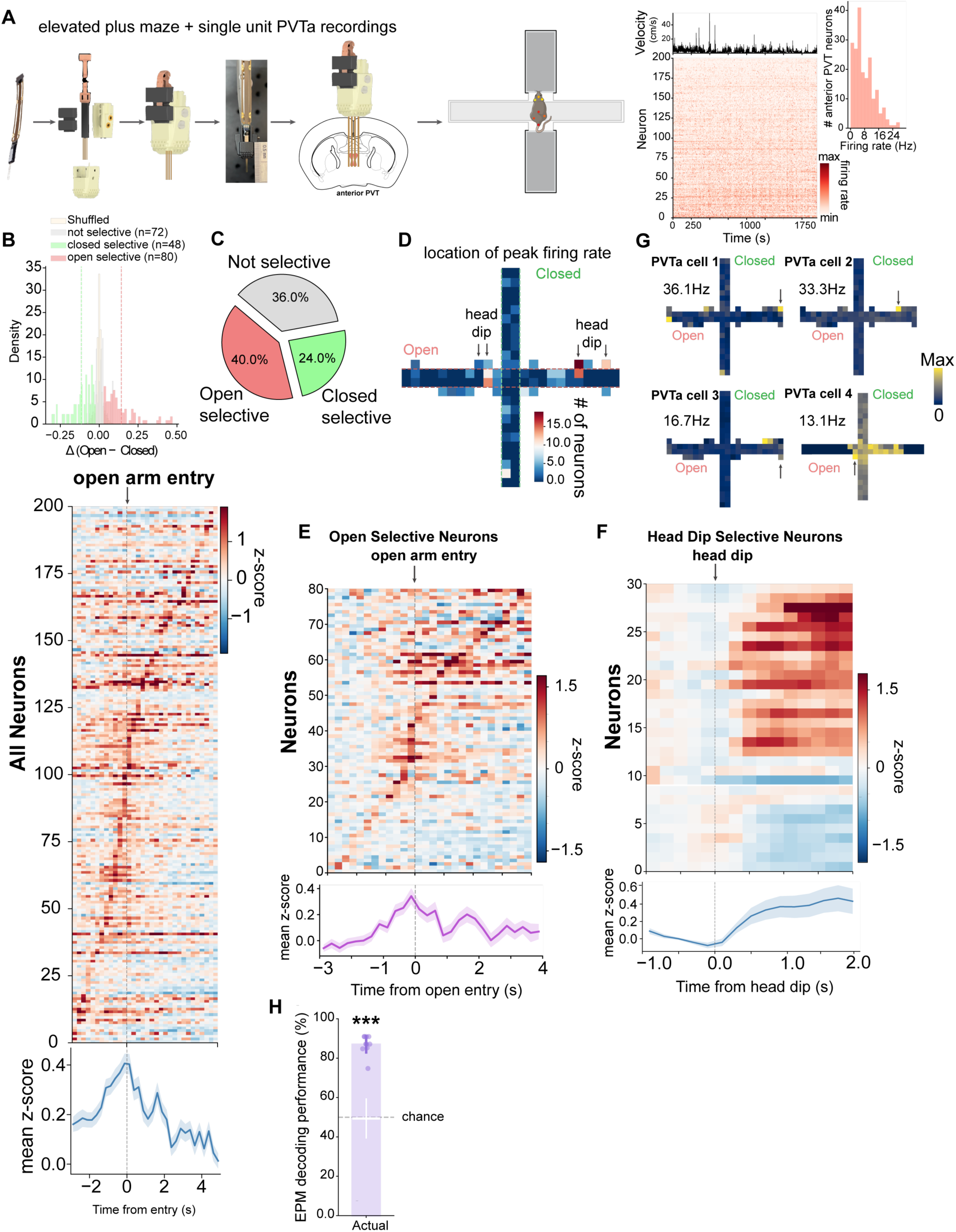
Coding of anxiety-related states in PVTa. **A**. Schematic of the design for chronic Neuropixels recordings into the PVTa. *Right panel* shows recording during EPM behavior with DLC behavioral tracking. Average velocity collected across mice across the entire session, *below* heatmap of all units recorded normalized to their max firing rate, *right* distribution of firing rates across all neurons. **B.** Single neural activity difference distribution between open and closed arms of the maze, with a breakdown in selectivity compared to shuffle, below heatmap of all neurons during open arm entry. **C**. Pie chart showing the percentage of neurons that are selective for each compartment of the maze. **D.** Spatial heat map of peak firing locations across the elevated plus maze, showing the distribution and density of neurons whose maximal firing occurs at each position, with the highest concentration in the head-dip zones. **E**. Heatmap of open arm selective units when animals transition into the open arm and average z-scored population activity. **F.** Heatmap of head dip-selective neurons when animals transition into the head dip zone, with the population average shown below. **G**. Example spatial heatmaps of individual neurons showing their peak firing rate in the maze with arrows pointing to head dip spatial bins. **H**. Decoding performance of a linear SVM trained on half the time bins when animals enter each zone versus shuffled data in white (*P* = 5.45 x 10^-8^). Data shown as mean ± SEM. Statistical significance determined by two-sided Mann-Whitney U test, where * *P* < 0.05, *** *P* < 0.001

### Aversive state representations in vSub depend on PVTa input

To test the functional role of the PVTa-to-vSub pathway, we used an intersectional approach to selectively manipulate vSub-projecting neurons in PVTa. We injected AAV2retro-Cre into vSub and a Cre-dependent inhibitory DREADD (AAV-DIO-hM4Di) or control vector into PVTa (Fig. 3A). In the elevated plus maze, CNO increased exploration of the center and open arms in DREADD-expressing mice (Fig. 3B), indicating reduced avoidance of anxiogenic regions. PVTa to vSub inhibition also increased center time in the open field (Fig. 3B), confirming that PVTa to vSub activity is required for normal avoidance behavior. Importantly, these changes in avoidance response were not accompanied by an overall increase in general locomotion, suggesting that inhibited mice actively chose to explore rather than engage in random hyperlocomotion (Supplemental Fig. 3E-F). To test the impact of acute excitation, we expressed ChRmine in PVTa neurons and optically stimulated their axons in vSub in a closed-loop fashion upon open-arm entry (Fig 3C). This stimulation likewise reduced avoidance, suggesting that imposing nonphysiological activity patterns onto this pathway disrupts the endogenous activity that supports avoidance and aversive state processing (Fig. 3D). This was further supported by doing 5 min OFF/ON open-loop stimulation in both the EPM and OFT tasks, showing a reduction in avoidance behavior during laser ON periods (Supplemental Fig. 3G-H).

**Figure 3:**
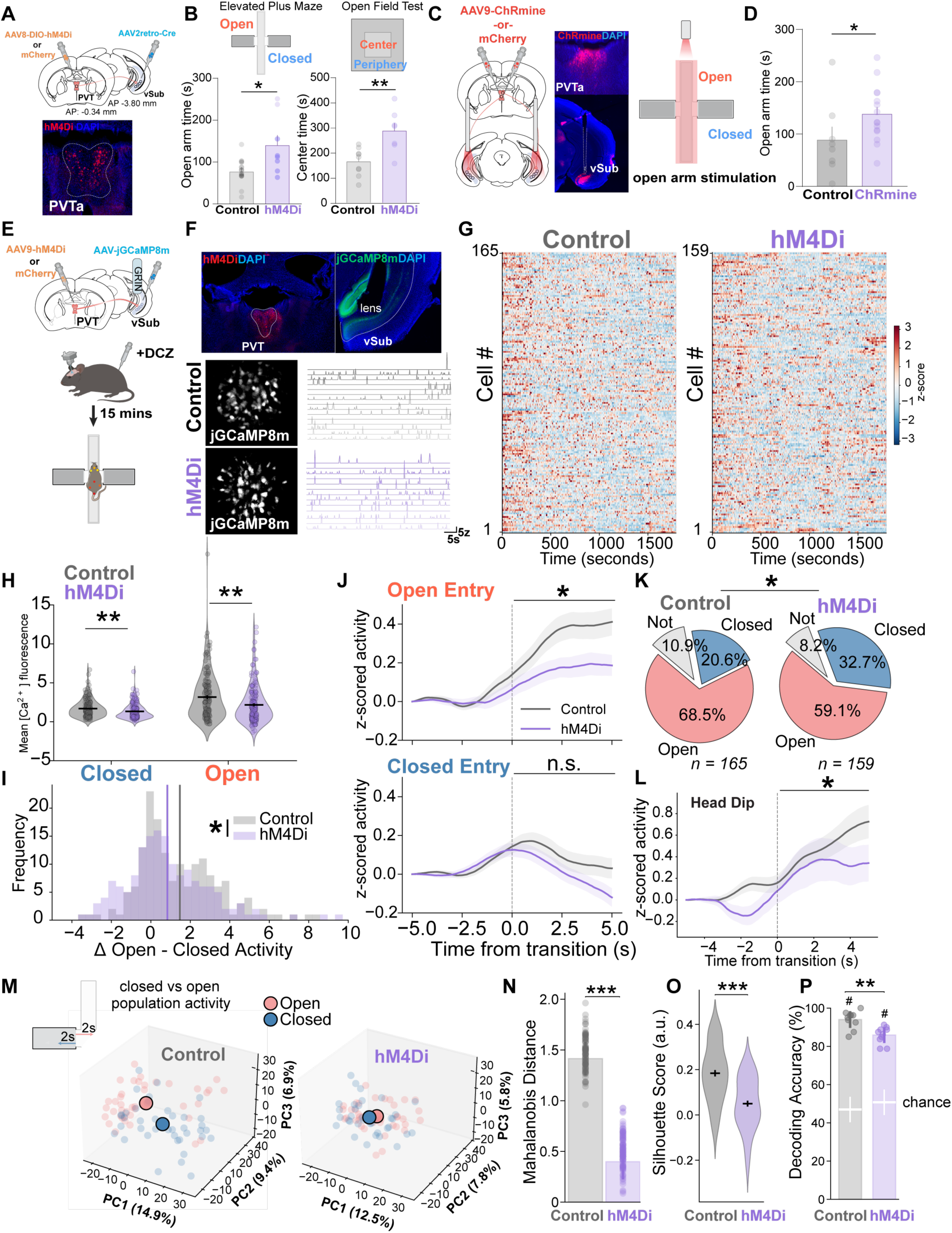
PVTa→vSub inhibition reduces avoidance behavior and weakens aversive-state representations in vSub. **A**. Design using conditional expression of inhibitory chemogenetics (hM4Di) in PVT^vHPC^ (*n* = 7-10 per group, male and female mice). **B.** EPM open arm time between each group (*P* = 0.043), *left panel*. OFT center time spent (*P* = 0.0052), *right panel*. **C**. Optogenetic closed-loop experiment using red-shifted excitatory opsin ChRmine, expression in the PVTa and bilateral optic fibers placed over vSub schematic and histology. **D.** Time spent in the open arms of the maze *right panel* (*P* = 0.047). **E**. Design schematic for inhibitory chemogenetic silencing (hM4Di) in PVTa during vSub imaging through a GRIN lens. Deschloroclozapine (DCZ) was administered 15 minutes before elevated plus maze testing. **F.** Expression of hM4Di in PVTa and jGCaMP8m expression in vSub of an example mouse. *Bottom panels*. Example FOVs from each group and extracted calcium events using. **G.** Normalized neural activity across all animals for the entire session (Control *n* = 165 cells/8 mice, hM4Di *n* = 159 cells/9 mice). **H.** Mean calcium fluorescence for both groups, closed arm (*P* = 0.000818), and open arm (*P* = 0.000716). **I.** Distribution of differences in single neural activity from open and closed arms (*P* = 0.0165) with vertical lines depicting population averages for each group. **J.** Time-locked activity when animals transition into either the open arm (*top panel*) and closed arms (*bottom panel*), with a significant difference when comparing AUC of the 5 seconds post transition between groups in open transitions (*P* = 0.0271) but not closed transitions (*P* = 0.172). **K.** Number of open and closed selective neurons (χ^2^ = 6.201, *P* = 0.0448). **L.** Mean time-locked activity during head dips (*P* = 0.0447). **M.** PCA analysis plotting the first 3 components for population activity in the first 2 seconds of entry into a zone, means shown with larger circles. **N.** Mahalanobis distance calculated between closed and open arms in the first 3 PCs (*P* = 2.56 x 10^-34^). **O.** Silhouette scores showing less separable closed and center clusters in the hM4Di group (*P* = 2.97 x 10^-9^). **P.** Linear SVM decoding results between open and closed zones between groups (*P* = 0.00355), shuffle shown in white (Control vs. shuffle *P* = 0.000181, hM4Di vs. shuffle *P* = 0.000180). Data shown as mean ± SEM. Statistical significance determined by two-sided Mann-Whitney U test except for panel K uses a χ^2^ test; * *P* < 0.05, ** *P* < 0.01, *** *P* < 0.001. # signifies *P* < 0.05 comparing real data to shuffled.

To assess how PVTa input modulates anxiety-related coding in vSub, we combined unilateral chemogenetic inhibition of PVTa with *in vivo* calcium imaging in vSub to observe changes in neural dynamics while minimizing behavioral changes. We expressed hM4Di in PVTa neurons (hM4Di) and GCaMP8m in vSub, implanted a GRIN lens over vSub, and performed single-cell microendoscopic imaging with miniaturized microscopes as mice explored the elevated plus maze (Fig. 3E-G). Relative to control mice, PVTa inhibition significantly reduced open-arm evoked vSub activity, indicating diminished responses to anxiogenic portions of the maze (Fig. 3H, Supplemental Fig. 3I). Quantifying each neuron’s differential activity in open versus closed arms, we found fewer neurons that discriminated between these compartments in hM4Di mice compared to controls (Fig. 3I). Analysis of vSub activity during distinct trajectories into the open and closed arms showed that, in control mice, vSub activity increased as animals approached and explored the open arms, whereas this recruitment was markedly blunted in hM4Di mice; activity suppression upon entry into closed arms was similar across groups (Fig. 3J). Single-cell selectivity analysis corroborated these findings: in the hM4Di group, fewer vSub neurons were open-arm selective (59.1 % vs. 68.5 %), and more were closed-arm responsive (32.7 % vs. 20.6 %) (Fig. 3K). Consistent with our PVTa recordings, in which head dipping at the open-arm edge strongly activated PVTa neurons, inhibiting PVTa with hM4Di reduced vSub responses to head dips, suggesting that these PVTa-dependent vSub signals, preferentially engaged in high-threat positions and during risk assessment, convey an aversive internal state to vSub rather than simply encoding maze location (Fig. 3L).

Dimensionality reduction and visualization of population activity during approach to open arms versus retreat to closed arms revealed that these population responses were more overlapping and less separable in hM4Di mice than in controls (Fig. 3M-O), and a classifier trained on vSub population activity showed reduced accuracy in distinguishing approach versus retreat in the hM4Di group (Fig. 3P). Together, these results demonstrate that PVTa input is required both for appropriate avoidance of anxiogenic environments and for vSub population codes that represent an aversive state and distinguish threat from safety.

### PVTa shapes vSub responses to motivationally salient stimuli

These findings indicate that PVTa input is necessary for vSub to encode ongoing threat during exploration of an anxiogenic environment. We next asked whether PVTa similarly configures vSub population structure after associative learning, when aversive predictors and outcomes can reweight vSub coding toward threat-related information. To test this, mice were trained in a trace conditioning task, where one odor predicted sucrose reward delivery (reward CS+), another predicted a brief aversive tail shock (shock CS+) and another was unreinforced (CS-) (Fig. 4A, Supplementary Fig. 3A). After training, mice learned to lick in anticipation of the reward CS+ but not the shock CS+ or CS-(Fig. 4B). To test how unilateral silencing of PVTa affected vSub encoding of conditioned stimuli (CS) and outcomes we performed two-photon imaging of GCaMP8m in vSub in control mice and in mice in which we inhibited PVTa neurons with hM4Di (Fig. 4C-D). Comparing control mice with mice in which PVTa was inhibited, we found a modest reduction in vSub responses to the rewarded CS+, and a reduction to aversive CS+ cues, suggesting that PVTa inhibition produces a cue-general reduction in vSub encoding of conditioned stimuli (Fig. 4D, G, and Supplementary Fig. 3B). In contrast, the responses to unconditioned stimuli (US) showed a striking pattern: PVTa-inhibited mice exhibited enhanced vSub responses to the rewarded sucrose US, but reduced responses to the aversive shock US (Fig 4D-E). We corroborated these results using feature-responsivity metrics, where we found that PVTa inhibition increased the fraction of vSub neurons responsive for reward delivery and the trace interval, and reduced the fraction responsive to shock (Fig. 4F). Supporting this, population decoding with a linear support vector machine classifier, PVTa inhibition decreased discriminability between the rewarded CS+ and neutral CS–, and between the shock CS+ and neutral CS–, indicating weakened encoding of conditioned cues (Fig. 4H-I, Supplementary Fig. 3D). By contrast, discriminability increased for the rewarded US relative to the time-matched post-CS− interval and decreased for the shock US (Fig. 4H–I, Supplementary Fig. 3D); consistent with the single-cell effects. Together, these results indicate that PVTa input provides an aversive-state control signal that biases vSub representations toward threat-predictive cues and aversive outcomes, while constraining reward-related activity.

**Figure 4:**
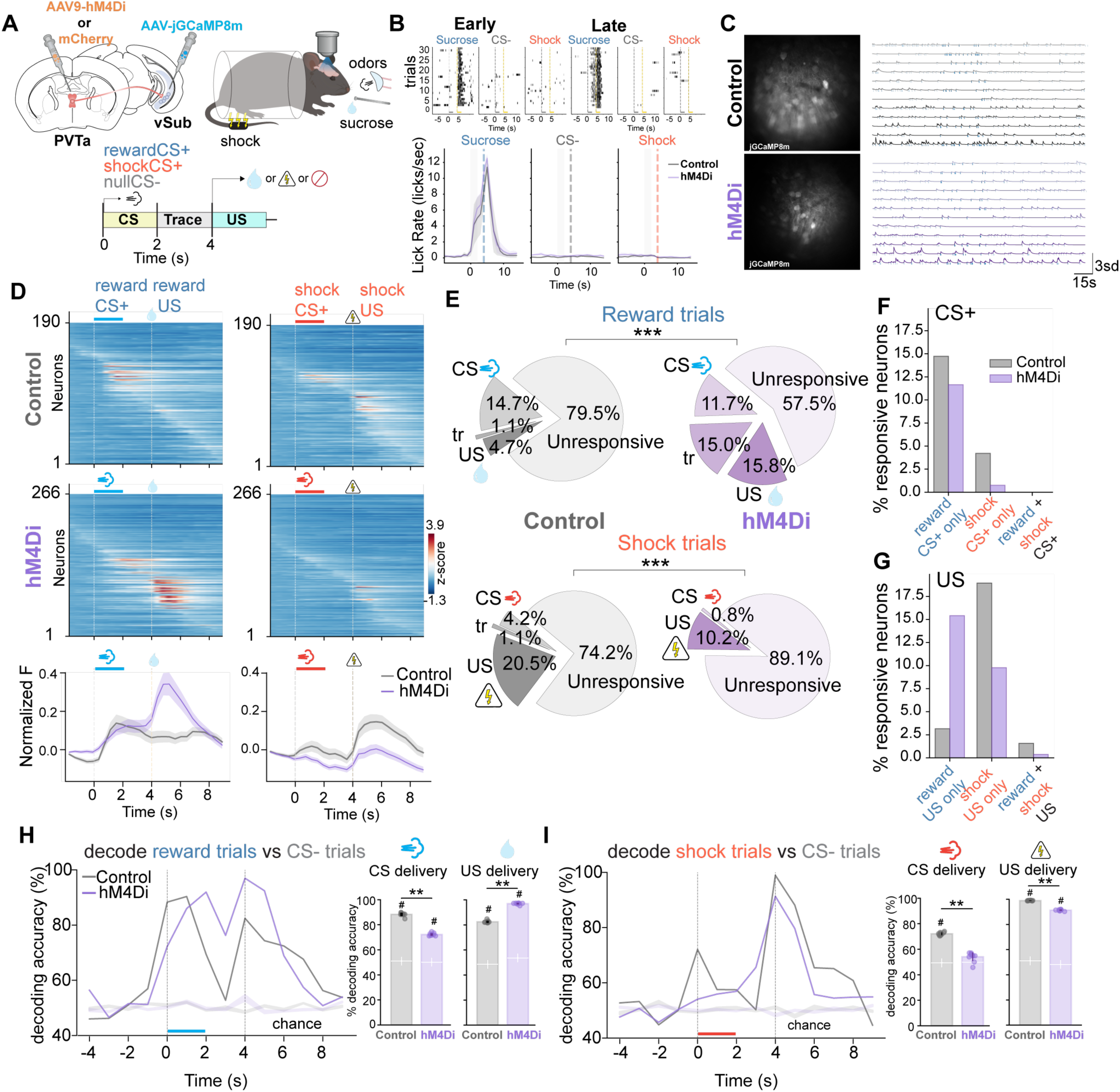
PVTa biases vSub coding toward threat over reward. **A.** Schematic of the odor-outcome associative learning task. Mice were trained to associate three distinct neutral odors with different outcomes: sucrose reward, no outcome (CS–), or mild tail shock, with odors presented for 2 seconds followed by a 2-second trace interval before US presentation. **B.** Example mouse lick raster plots from control animals during early and late stages of learning, and group-level average lick rates late in learning for each trial type. **C.** Example 2-photon imaging FOVs and extracted calcium traces recorded from vSub in each group. **D.** Heatmaps of z-scored calcium transients across neurons for reward and shock trials, with corresponding population averages activity shown on the bottom. **E.** *Top:* Proportion of neurons classified as CS-responsive, trace-responsive or US-responsive during sucrose trials (χ^2^ = 44.5, *P* = 1.2 x 10^-9^). *Bottom.* Proportion of neurons classified as CS-responsive, trace-responsive, or US-responsive during shock trials (χ^2^ = 20.1, *P* = 0.000165). **F-G.** Percentage of responsive neurons during either the CS period (*top panel*) and their overlap, and for the US period (**G**). **H.** Time-bin decoding accuracy for sucrose versus CS– between groups, with right panel bar charts showing the CS (Control vs. hM4Di *P* = 0.00586, Control vs. shuffle *P* = 0.00586, hM4Di vs. shuffle *P* = 0.00586) and US (Control vs. hM4Di *P* = 0.00582, Control vs. shuffle, *P* = 0.00586, hM4Di vs. shuffle *P =* 0.00582) decoding accuracy between groups using 1-second time bins post-onset. **I.** Same, but for shock versus CS–trials for CS (Control vs. hM4Di *P* = 0.00586, Control vs. shuffle *P* = 0.00586, hM4Di vs. shuffle *P* = 0.362) and US (Control and hM4Di *P* = 0.00582, Control vs. shuffle *P* = 0.00582, hM4Di vs. shuffle 0.00586). Data shown as mean ± SEM. Statistical significance determined by two-sided Mann-Whitney U test with Bonferroni correction; * *P* < 0.05, ** *P* < 0.01, *** *P* < 0.001. # signifies *P* < 0.05 comparing real data to shuffled.

## Discussion

Here, we have shown that the PVTa input to vSub implements a state-dependent gating mechanism that shifts hippocampal processing between threat and reward. Using viral tracing and barcoded single-cell anatomical approaches we found that a population of PVTa neurons that send a direct projection to vSub and previously unidentified bifurcating neurons to important limbic nodes. Using high-density Neuropixels recordings in the EPM, we found that PVTa population activity was enriched for neurons that preferentially fired in the anxiogenic open arms, with distinct subsets recruited at open-arm entry and during head-dip risk assessment, indicating that PVTa encodes aversive-space engagement and risk-assessment state at single-cell resolution. Further, we found that chemogenetic inhibition of the PVTa to vSub projection is required for normal avoidance of anxiogenic environments and for vSub population codes that encode an aversive state and reliably discriminate threat from safety. In a trace-conditioning task with vSub two-photon imaging, inhibition of PVTa reduced vSub coding of both reward-and shock-predictive cues, enhanced reward outcome responses, and diminished shock outcome responses, consistent with PVTa providing a state-dependent signal that biases vSub’s weighting of motivationally relevant information. Together, these data indicate that PVTa to vSub input provides a state-dependent control signal that maintains the balance of threat- and reward-related representations in vSub, prioritizing threat processing and cue salience during aversive states, and, when removed, diminishes threat discrimination with a relative enhancement of reward-related encoding.

Compared with previous results investigating the number of collateralizing PVTa neurons to the NAc, CeA, and BNST, the results are roughly similar. Overall, previous work specifically examining PVTa found that the highest percentage of dual-projecting neurons across any combination of downstream areas was ∼16%, ranging from 1% to 16^33,49,50^. One study reported that among ventral medial NAc shell-and dorsolateral BNST-projecting neurons, ∼15% are dual-projecting, which is similar to what we find in our study, where ∼17% are dual-projecting. Another study that examined the numbers of mPFC and vHPC collaterals across the entire anterior-posterior axis of the PVT, which found 1.6% bifurcation, our study identified 0.9%. Limitations of retrograde studies examining collateralizing PVTa neurons are similar in concept to MAPSeq, in that infection efficiency, targeting, quantification, and sampling of the downstream areas and PVT make direct comparisons difficult. In addition, while MAPSeq may remove low-sparsely projecting neurons, it has the advantage of assaying more than 3 projections, whereas previous studies have been limited to 2-3 targets and, by definition, represent a smaller subset of the neurons we find in our MAPSeq study. Further work will be needed to obtain accurate numbers, which is technically very challenging, as any technique suffers from the same issues of subsampling or high-throughput. We believe that while there may be disagreements in collateralizing neurons in our study compared to previous studies, our results expand what was previously known about PVTa neurons beyond its projections to NAc, CeA, and BNST, of which there is more than 1 study. In addition, the previous studies mentioned here were done in rats; whether there are species differences between mice and rats is unknown. Future studies can delineate the functional consequences of these collaterals and their relative abundance. While our manipulation studies using chemogenetics and optogenetics targeted PVTa neurons projecting to vSub, our anatomy showed that the majority of PVTa^vSub^ neurons are single projectors; a smaller fraction have collaterals to other areas. Therefore, the effects we see on vSub coding may result from collaterals that, in turn, cause changes in vSub activity. Future work can begin to delineate these collateral projections from the single projectors. It is possible that the collaterals are efference copies sent to downstream areas to properly coordinate behavior.

vHPC is a central hub in the broader circuitry that supports emotional and motivated behavior, with distinct cell types, inputs, and projection-defined outputs coordinating diverse approach- and avoidance-related functions^1–4,7,18^. At the representational level, recent population recordings suggest that vCA1 ensembles robustly encode the identity and sensory features of salient stimuli, including anxiogenic contexts, rewarding outcomes, and social cues, while maintaining distinct, stable representations for individual stimuli rather than collapsing them into a single generalized value axis^4^. This organization raises a key question: how are stable vCA1 identity codes dynamically weighted to drive state-dependent behavior, particularly during approach–avoidance decisions? Recent work has begun to define how upstream inputs, including from the amygdala^13–15^, convey aversive and appetitive information to the vHPC and shape its engagement during learning and anxiety-related behavior. Here, we identify PVTa as a previously underappreciated node in this circuit and show that it provides an aversive-state control signal that modulates vSub output, increasing the weighting of threat-related information in vSub population codes, and thereby promoting avoidance-related signaling in downstream targets and the formation of emotionally salient memories.

The PVT has been shown to be an integrative node that combines homeostatic and internal-state signals from hypothalamic and brainstem sources with information from limbic and cortical circuits to shape adaptive behavior^23,24,39,51–54^. Recent work supports functional specialization along the anterior–posterior axis of the PVT, with subpopulations in posterior PVT (PVTp) versus PVTa showing distinct population response profiles in bulk fiber photometry^55,56^. In a motivational conflict task, both PVTa and PVTp exhibited responses to conditioned cues and footshock, but only PVTp was robustly responsive to sucrose reward, indicating a bias in how appetitive outcomes are represented across the axis^55^. These populations have also been subdivided using genetic markers, suggesting that PVTp-biased subpopulations show strong responses to aversive stimuli with suppression during rewarding events, whereas PVTa-biased subpopulations are broadly inhibited by salient, arousing events^55^. Others have shown that PVTa is inhibited only by reward itself and is excited by cues that predict aversive outcomes and aversive stimuli^57^. However, an important caveat is that fiber photometry cannot resolve how individual neurons contribute to these bulk dynamics, leaving open the question of how single-cell heterogeneity underlies the observed population patterns. Our single-unit recordings in PVTa directly address this limitation by identifying a subpopulation strongly engaged during active threat evaluation, with elevated firing in the open arms of the elevated plus maze and during head-dip risk assessment. Together, these findings support a model in which PVTa encodes a sustained threat-state signal that can be transmitted to vSub. This motivates further cell-type-resolved experiments to define which PVTa populations carry threat-state information and how these signals are transformed within vSub microcircuits and projection-defined output pathways.

Together, these findings support a framework in which PVTa state signals configure vSub output by modulating how representations of stimulus identity are expressed and read out by downstream circuits. An important implication is that vSub can preserve stable identity codes, while internal-state-dependent gain control via PVTa biases the routing and impact of those codes to shape behavioral output. Building on this framework, delineating the specific PVTa inputs and vSub mechanisms that implement state gating will help explain how hippocampal output is flexibly reconfigured across contexts, and how disruptions in this process may contribute to motivational and salience-related dysfunction in stress-related neuropsychiatric disorders.

---

**Supplemental Figure 1.**
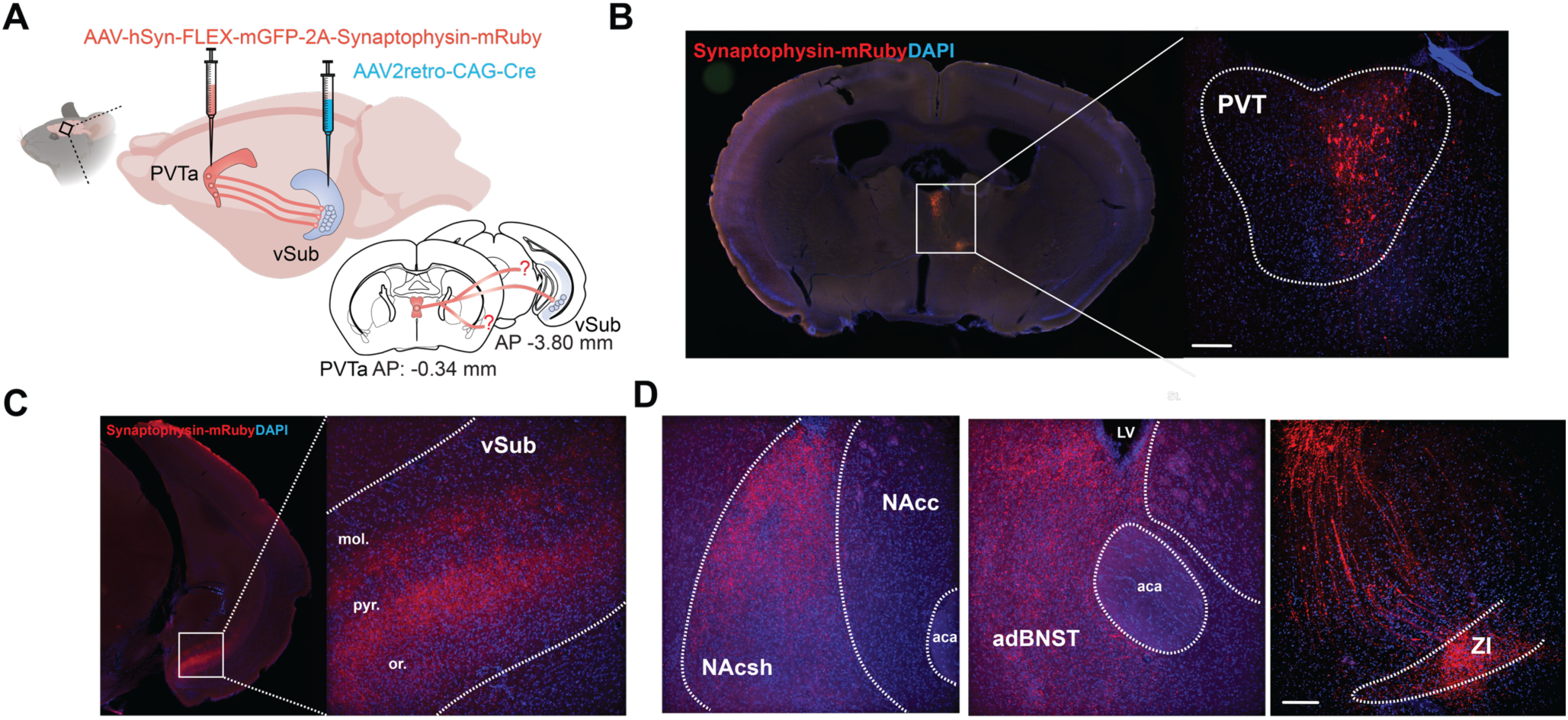
A. Design schematic for tracing collaterals of vHPC-projecting PVT neurons using conditional expression of Synaptophysin-mRuby in PVT. B. Coronal section of sample PVT cell bodies expressing mRuby with a zoomed image. C. Coronal sections showing PVT terminals labeled with mRuby in the hippocampus. D. mRuby expression within the NAc shell, BNST, and ZI.

**Supplemental Figure 2.**
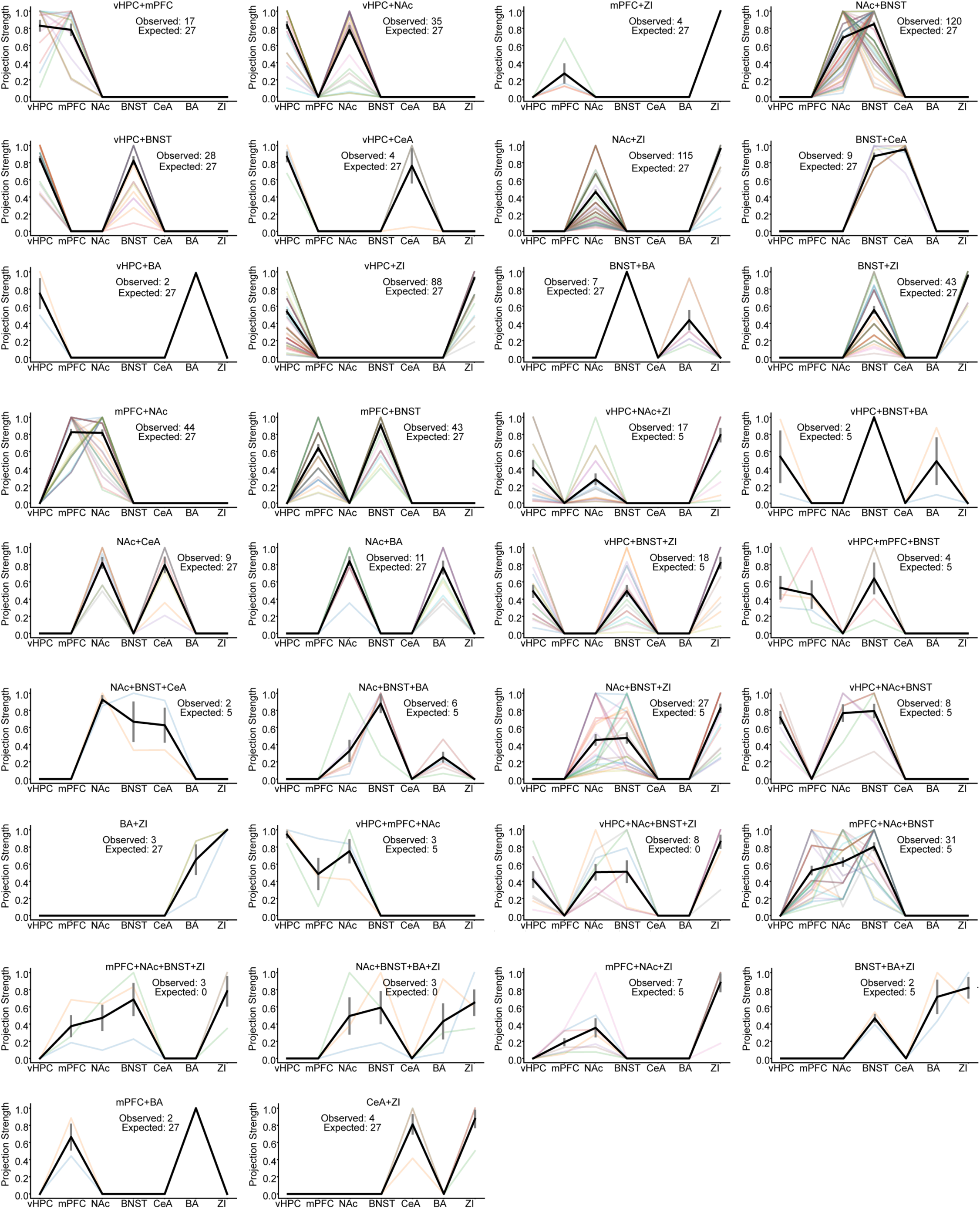
Plots of PVT projection motifs showing individual putative neuron projection strengths in different colors, with the mean of all neurons in black. These plots highlight the heterogeneous projections strengths within a motifs.

**Supplemental Figure 3.**
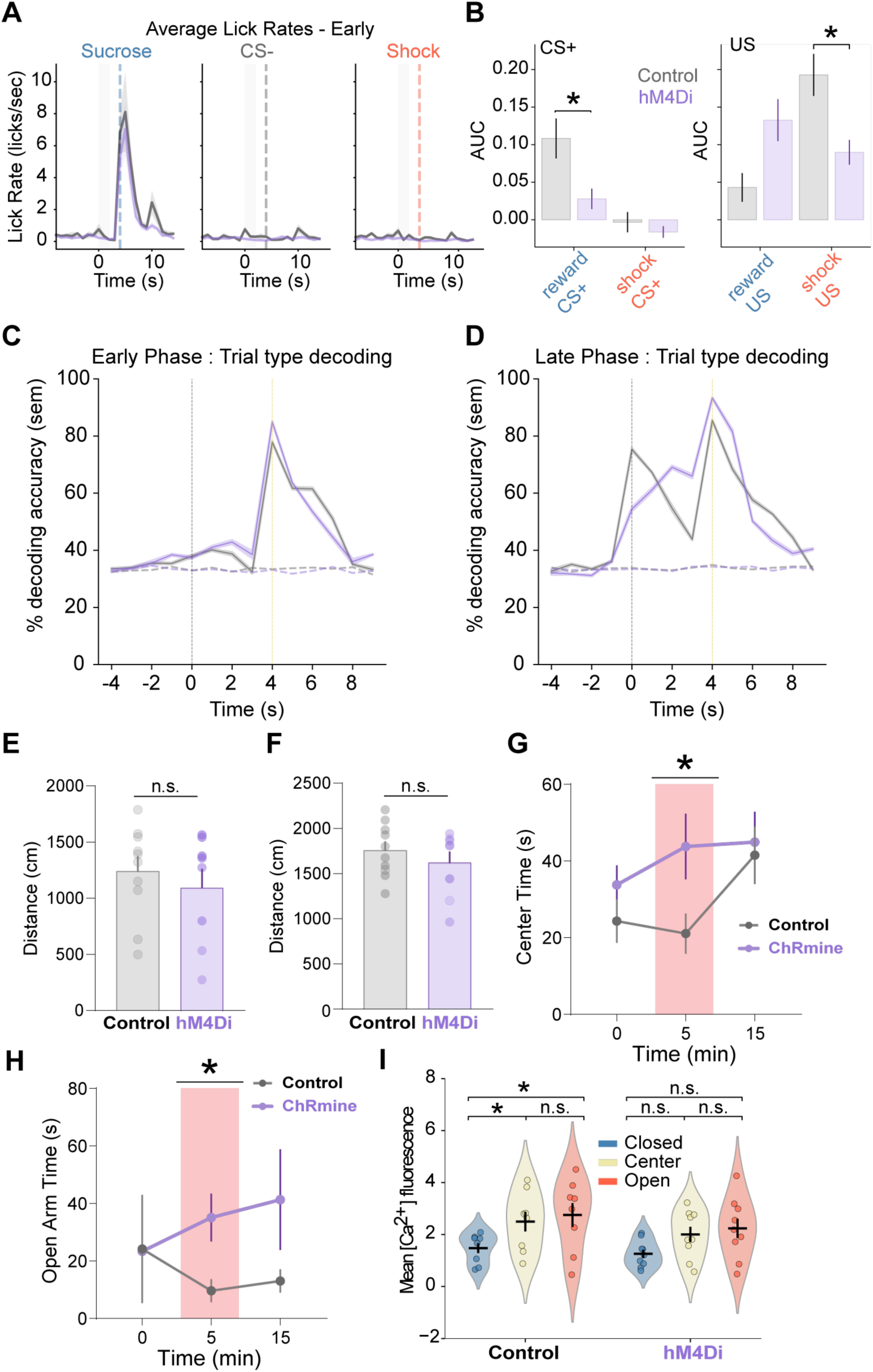
A. Early-session average lick rates across animals, by trial type. B. Mean population calcium event rates for each trial type during late sessions. C. Trial type decoding across time bins for the early session, D. and for the late session. E. Total distance in the elevated plus maze between controls and hM4Di animals for Fig 3B, left panel (*P* = 0.515). F. Total distance in the open field test between groups for Fig 3B, right panel (*P* = 0.536). G. Open-loop optogenetic excitation for 5min OFF/ON/OFF epochs in the open field test between groups (post hoc test with Bonferroni correction, *P* = 0.0173). H. Open arm time in the elevated plus maze after open-loop optogenetic excitation for 5 min OFF/ON/OFF epochs (post hoc test with Bonferroni correction, *P* = 0.0355). I. Subject-level average calcium activity in each compartment of the elevated plus maze within groups compared with Wilocxon-signed rank test and corrected with Bonferroni; for Controls closed vs. center (*P* = 0.023), center vs. open (*P* = 0.75), and closed vs. open (*P* = 0.047), all hM4Di comparisons (*P* > 0.05). Data shown as mean ± SEM. Statistical significance determined by two-sided Mann-Whitney U test unless otherwise stated; * *P* < 0.05, ** *P* < 0.01, *** *P* < 0.001.

## Acknowledgments

M.M.G. was supported by the National Institute of Mental Health (F31 MH130127) and the National Institute of Neurological Disorders and Stroke (DSPAN F99/K00 NS130927). M.A.K. was supported by the National Institute of Mental Health (R01 MH108623, R01 MH111754, R01 MH117961 and R01 MH125515), the National Institute on Deafness and Other Communication Disorders (R01 DC019813), the One Mind Rising Star Award, the Human Frontier Science Program (RGY0072/2019), the Esther A. and Joseph Klingenstein Fund, the Pew Charitable Trusts, the McKnight Memory and Cognitive Disorders Award, and the Ray and Dagmar Dolby Family Fund.

## Author contributions

M.M.G. and M.A.K. conceptualized the project. M.M.G. and M.A.K. were responsible for the methodology. M.M.G., G.T., K.L., M.L., A.K., J.S.B., S.H., J.T., and V.S.T. were responsible for the investigation. M.M.G. was responsible for visualization. M.M.G. wrote the first draft of the manuscript. M.M.G. and M.A.K. reviewed and edited the final version of the article.

## Methods

All procedures were conducted in accordance with the U.S. NIH Guide for the Care and Use of Laboratory Animals and the Institutional Animal Care and Use Committees at UCSF. Male and female adult C57BL/6J mice were supplied by Jackson Laboratory and were used beginning at 8-12 weeks of age. Mice were co-housed with littermates (2-5 per cage) in a temperature (22-24°C) and humidity (40-60%) controlled environment. Mice were maintained with unrestricted access to food and water on a 12-hour light/dark cycle, with tissue processed during the light phase. All subjects were randomly assigned to experimental conditions, with approximately equal numbers of male and female mice.

### General Stereotaxic Surgical Procedures

Mice were anesthetized with 1.5% isoflurane with an oxygen flow rate of ∼1 L / min, and head-fixed in a stereotactic frame (David Kopf, Tujunga, CA). Eyes were lubricated with an ophthalmic ointment, and body temperature was maintained at 34-37°C with a warm water re-circulator (Stryker, Kalamazoo, MI). Fur was shaved and incision site sterilized with isopropyl alcohol x3 and betadine solution x3 prior to beginning surgical procedures. Lidocaine HCl 2% solution was injected subcutaneously local to incision, and post-surgical analgesia was provided by meloxicam and slow-release buprenorphine. A craniotomy was made at injection site with a round 0.5 mm drill bit (David Kopf, Tujunga, CA). A Nanoject III syringe (Drummond Scientific, Broomall, PA) was used with a pulled glass pipette (tip width 20-30 µm) to inject viruses at a speed of 10 nl/sec described below. Targeting and injection parameters used for vSub were M/L ±3.0 mm and A/P −3.8 mm, 125 nl of virus was injected at a depth of −4.75 mm, 150 nl at −4.50 mm, 125nl at −4.25 mm, and 100nl at −4.0 mm (500nl total) with respect to bregma skull, and for PVT M/L − 0.05 mm, A/P −0.34 mm, and 350 nl injected at a depth of D/V −3.5 mm with respect to bregma skull.

### Synaptophysin tracing

Mice were first injected with AAV2retro-CAG-cre (4.1 x 10^12^ vg/ml, UNC Vector Core) and mixed with 1:5 Fluoromax beads for its total dilution of 1:50 (Thermofisher) in the left hemisphere of vSub. The needle was held in place for >1 minute prior to moving to the next D/V coordinate and remained in place for >5 minutes following the final injection before slowly removing from the brain. 3-5 days later, mice were injected with AAVDJ-hSyn-FLEX-mGFP-2A-Synaptophysin-mRuby (5.8 x 10^12^ vg/ml, Stanford vector core) in the left hemisphere of PVT. The needle was held in place for >5 minutes after the injection before being slowly removed from the brain. 3-4 weeks after the second injection surgery, mice were injected with a lethal dose of 2:1 ketamine/xylazine solution intraperitoneally. Mice were perfused transcardially using 4% PFA solution. Brains were extracted and placed in 4% PFA solution for 2-3 days to allow further fixation. After saturating with a 30% sucrose solution, coronal slices of 50 µm width were collected using a Leica SM2000 microtome. Slices were collected in a 1X PBS solution with 0.02% sodium azide, mounted onto glass slides, and coverslipped with Fluoromount G with DAPI (Southern Biotech, Birmingham, AL). Stitched images of full tissue sections were taken at 10X magnification using a BZ-X810 fluorescence microscope (Keyence, Itasca, IL). 10X and 20X images of areas of synaptophysin expression were taken at a Nikon Ti2-E microscope equipped with a CREST X-Light V2 large field of view spinning disk confocal. Images were then processed using ImageJ.

### MAPseq

#### Sample Generation

Experiments were conducted as previously described^18,46,47^. Adult male and female mice (n=9) were anesthetized and injected with 250 nl MAPseq Sindbis viral barcode library (3×10^10^ GC/ml, diversity of 2×10^7^ different barcode sequences, Cold Spring Harbor Laboratories) into the PVT. The titer and volume of Sindbis virus used are like that previously published, in which single-cell analysis indicated that on average, each neuron expressed one barcode, with a small fraction expressing more than one^46^. However, as discussed at length previously^46–48^, if neurons express more than one barcode, this would not change the distribution of projection patterns of assayed neurons. While the total number of traced neurons would be overestimated, the relative abundance of each projection motif type and bulk connection strength would not be changed. We predict that >99% of cells would be labeled uniquely if the fraction of unique labeled cells F=(1-(1/N))^k-1^ where N is the barcode diversity and k is the number of infected neurons^46^. ∼44-46 hours after surgery, mice were rapidly decapitated, and brains were flash-frozen in a slurry of 2-methylbutane (Fisher Scientific) on dry ice and stored at −80° C until processing. Brains were then embedded in O.C.T. (Fisher Scientific) and sectioned coronally on an HM525 cryostat (Fisher Scientific) with 100-200 µm thickness of target areas with a new blade between each region: vSub, mPFC, NAc, BNST, ZI, BA, and CeA, negative control area (DLS) and the source region of interest (PVT) were mounted on SuperFrostPlus (Fisher Scientific) slides and kept on dry ice. Brain punches of target areas were collected ipsilateral to injection site (while avoiding fiber tracts) from good quality consecutive sections, in which only 1 area was punched from any individual section to minimize any potential contamination, on dry ice with a chilled 500 µm puncher (Electron Microscopy Instruments), which was cleaned between each brain area with 100% ethanol, and samples were stored in 1.5 ml microcentrifuge tubes from each mouse and kept on dry ice until processing. Each sample was then homogenized in 400 µL of TRIzol (Thermofisher) and vortexed, followed by a quick spin, and then kept on dry ice before shipping to MAPseq Core Facility for barcode extraction and sequencing (Cold Spring Harbor Laboratories).

Barcode extraction and sequencing were previously described^18,46,47^. In brief, total RNA was extracted from each sample using TRIzol reagent (Thermo Fisher) and sample RNA with spike-in RNA was mixed (obtained by in vitro transcription of a double-stranded ultramer with sequence 5′-GTCATGATCATAATACGACTCACTATAGGGGACGAGCTGTACAAGTAAACGCGTAATGATACGGCGACCACCGAGATCTACACTCTTTCCCTACACGACGCTCTTCCGATCTNNNNNNNNNNNNNNNNNNNNNNNNATCAGTCATCGGAGCGGCCGCTACCTAATTGCCGTCGTGAGGTACGACCACCGCTAGCTGTA CA-3′ (IDT)30) and reverse transcribed using a gene specific primer 5′-CTTGGCACCCGAGAATTCCANNNNNNNNNNNNXXXXXXXXTGTACAGCTAGCGGTGGTCG-3′, where X is one of >300 true-seq-like sample-specific identifiers and N12 is the unique molecular identifier, and SuperscriptIV Reverse Transcriptase (Thermo Fisher, with manufacturer instructions). All first-strand cDNAs were pooled and purified using SPRI beads (Beckman Coulter) to produce double-stranded cDNA. Samples were treated with ExonucleaseI (NEB) and performed two rounds of nested PCR using primers 5′-CTGTACAAGTAAACGCGTAATG-3′ and 5′-CAAGCAGAAGACGGCATACGAGATCGTGATGTGACTGGAGTTCCTTGGCACCCGAGAATTCCA-3′ for the first PCR and primers 5′-AATGATACGGCGACCACCGA-3′ and 5′-CAAGCAGAAGACGGCATACGA-3′ for the second PCR using Accuprime Pfx polymerase (Thermo Fisher). Finally, PCR amplicons were gel extracted using Qiagen MinElute Gel extraction kit and library sequenced on an Illumina NextSeq500, set at high-output run and paired-end 36, using the SBS3T sequencing primer for paired-end 1 and the Illumina small RNA sequencing primer 2 for paired-end 2.

All preprocessing of sequencing data was performed at the MAPseq Core Facility at Cold Spring Harbor Laboratories exactly as described in Kebshull 2016 (Supplementary note 4)^46^. All sequencing was done blinded to sample identity. Briefly, Illumina sequencing results (.fastq files) were merged into one file that contained paired end 1 (barcode sequence) and paired end 2 (the 12-nt unique molecular identifier (UMI) and 8-nt slice-specific identifier (SSI)), so that each line contained corresponded to a single read containing the 30-nt barcode, the 2-nt pyrimidine anchor (YY), the 12-nt UMI and the 8-nt SSI. Reads were de-multiplexed based on SSI using, fastx_barcode_splitter tool (http://hannonlab.cshl.edu/fastx_toolkit/commandline.html#fastx_barcode_splitter_usage) and filtered to remove ambiguous bases, and collapsed to unique sequences and sorted. Next, a threshold was selected for how many reads a sequence must be used in the analysis. As in Kebshull et. al 2016^46^, a minimum read threshold was manually selected to remove the long tail of the sequence rank profile of the Illumina results so as to avoid contamination with PCR and sequencing errors. The remaining reads were collapsed after the removal of the 12-nt of the UMI to convert reads into counts. Spike-in molecules, 24-nt barcodes followed by the sequence ATCAGTCA, were split from the virally expressed barcodes and processed separately. Error correction was performed using the short-read aligner *bowtie*, which generated all possible alignments of barcode sequences (>1 count, allowing up to 3 mismatches). All barcode sequences that mapped to each other were found, and low-complexity sequences were removed by filtering barcodes with stretches of more than 6 identical nucleotides.

Raw barcode reads were then normalized by a relative number of spike-in RNAs and organized into an N x R matrix with N barcodes detected in R possible regions. The N barcodes are a proxy for the number of cells sending projections in the source region; in the text, we refer to each row as a cell. Each value in a row indicates the number (non-negative integer values) of detected barcodes in that region, which corresponds to the strength of the projection (density of axons) to that region. Matrices were concatenated for all mice to obtain a single larger matrix. To limit analysis only to cells that project to at least one region, we removed rows that contained all zeros. We threshold-filtered data based on the presence of 10-fold enrichment of source (PVT) barcode reads compared to at least one target area and the removal of any barcode with a read count in the negative target area (DLS) prior to further analyses. Lastly, to compare projection patterns between neurons on the same scale, we normalize each row in the N x R matrix to that row’s maximum value, such that raw barcode counts are rescaled between zero and one.

### Data analysis

#### Conditional probability

To calculate the conditional probability *P*(*B* | *A*), for a pair of regions A and B, we first found the number of cells that project to region A, inclusive (i.e. cells that may project to A alone or A with projections to other regions as well), denoted *N*_A_. We then found the subset of cells in this group that projects to region B, inclusive (i.e. cells that may project to A and B and may or may not have projections to additional regions), denoted *N*_B|A_. The conditional probability is the proportion of cells projecting to B within the subset of cells that project to A, i.e. 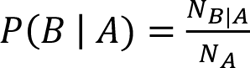

#### Over and under-represented motifs

To quantify the significantly over and under-represented projection motifs in our data, we developed a null model with which to compare. First, we note that our data is a matrix of neurons by target regions, and when analyzing projection motifs, we first binarized this matrix to only consider whether a neuron projects to a particular target, regardless of its projection strength. This binarized matrix is then interpreted as a bipartite graph where the neurons form one node set and the target regions form the other node set and values of 1 in the matrix indicate an edge between nodes^58^.

We then constructed our null model as an Erdős–Rényi random (bipartite) graph where edge formation is determined by a constant probability following a binomial distribution^48,58^. This results in a null model where neurons are assumed to not have any intrinsic preference for projecting to target regions and where the probability of projecting to one region is conditionally independent from that of another region.

The probability of edge formation denoted *p_e_*, determines the edge density of the null model, which we assumed is equal to our empirical data. In generating a random graph using *p_e_*, some simulated neurons will have zero projections to any of the target regions, however, due to the nature of MAPseq, the empirical data only includes neurons that have at least one projection to the target regions sampled. To properly model our data, we need to know the total number of neurons, including those that had no projections to one of our target regions. We can estimate this by noting that *N_f_* = *N*_0_ ⋅ *p_e_*, where *N_f_* denotes the number of neurons in our empirical data and *N*_0_ refers to the total number of neurons, such that *N*_0_ − *N_f_* would be the number of neurons with zero projections^47^.

First, we infer *N*_0_ from the empirical data by assuming a binomial model and recalling that the probability of at least one projection is one minus the probability of no projections:

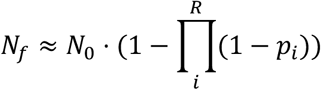

Where *p_i_* refers the probability of a neuron projecting to region *i* among the total *R* regions. We define 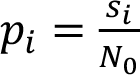 where *s_i_* is the number of neurons in the empirical data that project to region *i*. By substitution, we get:

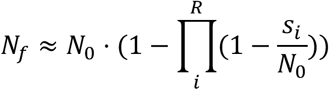

We now have a polynomial with one unknown, *N*_0_, and we can solve this using any root-solving algorithm. Knowing *N*_0_ and *N_f_*, we can then solve for *p_e_*:

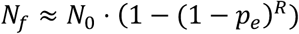

This is because the number of neurons with one at least projection using the null model with *p_e_* must reproduce the number of neurons in the empirical data which are also filtered to only include those with at least one projection. This equation is also a polynomial with one unknown, *p_e_*, and again, we can solve it numerically using a root-solving algorithm. In our data, *N*_f_ = 1888, and we computed *N*_0_ ≈ 2753, *p*_e_ ≈ 0.152. Using *p*_e_ and *N*_0_ we can compute the expected counts for each motif, e.g. the expected counts for each single-projector motif under the null model are computed as *N*_0_ ⋅ *p*_e_ ⋅ (1 – *p*_e_)^6^, since there are 7 target regions. We then compute the two-sided p-values for our observed counts under the null binomial model using the binomial test.

All MAPseq data was analyzed using custom scripts written in Python (available at github.com/mkheirbek).

### Freely moving behavior

#### Elevated Plus Maze

Mice were placed in a standard EPM sized maze (13.5’’ height of maze from floor, 25’’ full length of each arm type, 2’’ arm width, 7’’ tall, closed arms, with 0.5’’ tall/wide ledges on the open arms made by Maze Engineers, Skokie, IL), with ∼450 light lux centered over the open arms to promote avoidance. Mice were placed in the center region of the maze and were allowed to explore for 10 (manipulation experiments) or 30 (imaging experiments) minutes while recording behavior with a webcam EthoVision XT 10 (Noldus, Leesburg, VA).

For imaging experiments, behavior was tracked using DeepLabCut^59^ software (version 2.3.9). 8 positions (nose, head, left & right ear, left & right trunk, mid-trunk, base of tail), and used the body position to determine open, center and closed arm occupancy. We used a ResNet-50-based neural network with default parameters for (1003000 x) 2 training iterations. We used a p-cutoff of 0.95 to condition the X, Y coordinates for future analysis after validation then used custom code to further filter data for discontinuities.

#### Open Field Test

Mice were placed in an arena (18’’ x 12’’ length-width-height; Kinder Scientific, Poway, CA) with bright light (∼650 lux) centered over the center zone and allowed to explore for 15 minutes while behavior was recorded and analyzed with EthoVision XT 10 (Noldus, Leesburg, VA) software.

### Manipulation experiments

#### DREADDs

Mice were first injected with AAV2retro-CAG-Cre (4.1 x 10^12^ vg/ml, UNC Vector Core) bilaterally into vSub. The needle was held in place for >1 minute prior to moving to the next D/V coordinate and remained in place >5 minutes following the final injection before slowly removing from the brain. PVT was bilaterally injected with 350nl of either AAV8-hSyn-DIO-hM4Di(Gi)-mCherry (2.3 x 10^13^ vg/ml, Addgene #44362), or AAV8-hSyn-DIO-mCherry (2.5 x 10^13^ vg/ml, Addgene #50459). The needle was held in place >5 minutes after the injection before slowly removing from the brain. For behavioral experiments, animals were injected with 3mg/kg of clozapine-N-oxide (CNO, NIH) dissolved in dimethyl sulfoxide (Sigma-Aldrich) 15 minutes prior to behavior.

#### Optogenetics

PVT was bilaterally injected with 350nl per site with either AAV8-nEF-ChRmine-mScarlet (2.1 x 10^13^ vg/ml, Addgene #137158) or AAV9-hSyn-mCherry (2.4 x 10^13^ vg/ml, Addgene #114472). Following a ∼5 mm long 200 mm core, 0.37 numerical aperture (NA) multimode fiber (ThorLabs) with at least 70% light recovery was implanted into each hemisphere above vSub at M/L ±3.0 mm A/P −3.8 mm, and DV −4.25 mm from bregma skull. Two anchoring screws were placed and cemented down with metabond to secure optic fiber implants.

During tasks, 5 ms pulses of ∼8-10 mW at 20Hz light were delivered via a 565 nm 100 mW laser (Opto Engine, Midvale, UT) to fiber optics implanted in mouse brain using a fiber optic patch cable. Light delivery was controlled via TTL pulse delivered from EthoVision XT 10 and Noldus IO box system to a Master-8 stimulator (AMPI, Jerusalem, Israel) for closed-loop stimulation. The laser was only triggered ON when mice were tracked live with EthoVision in a pre-drawn stimulation zone; center and open arms for EPM.

#### Calcium imaging

PVT was injected with either AAV9-hSyn-hM4Di(Gi)-mCherry (2.6 x 10^13^ vg/ml, Addgene #50475) or AAV9-hSyn-mCherry (2.4 x 10^13^ vg/ml, Addgene #114472) and vSub was injection with AAV1-hSyn-jGCaMP8m-WRE (2.5 x 10^13^ vg/ml, Addgene #162375) diluted 1:1 with sterile 1x PBS. A needle track was created by lowering a 27’ gauge blunt needle into vSub DV −4.65 mm at 100 µm per second. Following either a 0.6 mm diameter x 7.3 mm length (Inscopix, #1050-004413) or 0.5 mm diameter x 6.1 mm height (Inscopix, #1050-004415) Proview Integrated GRIN lens was implanted at a speed of 10 µm per 5 seconds over vSub at ML −3.0 mm, AP −3.80 mm, and DV −4.70 mm from bregma skull. Following a skull screw and custom-made titanium headbar was then attached to the skull and cemented in place with GRIN lens using metabond. 15 minutes prior to behavior and neural recordings, mice were injected with 100 µg/kg deschloroclozapine (DCZ, HelloBio).

#### 2-photon imaging

Procedure and analysis were done as mentioned in previous work^3,4^. Images were captured using an Ultima IV laser scanning microscope (Bruker Nano, Middleton, WI, acquired with Prairie View 5.4) equipped with resonant scanning mirrors and high-speed scan electronic controller, dual GaAsP PMTs (Hamamatsu model 7422PA-40), and motorized z focus (100 nm step size). GCaMP signal was filtered through an ET-GFP (FITC/CY2) filter set. Laser signal was provided by a MaiTai DeepSee mode-locked Ti:Sapphire laser source (Spectra-Physics, Irvine, CA) providing > 150kW max output at 920 nm. Acquisition speed was 30 Hz for 512 x 512 pixel images. Images were averaged 8x offline, yielding a final frame rate of 2.5 Hz. Prior to each conditioning session, 2 imaging fields of view (FOVs) were determined, and imaging was conducted at those FOVs for the entire session using an objective z-piezo stage (Bruker, Middleton, WI) that rapidly (∼80 ms) switched between different z-planes. Each FOV was separated by > 100 µm in the z-dimension (dorsal-ventral) to ensure no overlap of cells across different FOVs. To facilitate the re-identification of a specific FOV across sessions, the top of the GRIN lens served as a reference z-plane. Optimal laser power was determined for each FOV based on GCaMP expression level and was kept constant across sessions for a specific FOV. For each trial, imaging began 5 seconds prior to stimulus onset and was terminated 6 seconds after the US presentation.

Videos were motion corrected offline using non-rigid motion correction based on template matching (Suite2p (Pachitariu et al., 2017)). Cell segmentation and calcium transient time series data were extracted using CNMFe. Putative neurons were manually inspected for appropriate spatial properties and Ca^2+^ dynamics and were visually checked against the corresponding motion-corrected video in ImageJ. Ca^2+^ transient events were extracted using the OASIS algorithm (Friedrich et al., 2017) embedded within CNMFe. We used these inferred calcium events for all analyses unless otherwise noted. Denoising (CNMFe) and deconvolution (OASIS) steps were applied.

#### Associative Learning

Animals were water restricted to ∼85-90% ad lib weight during experiments. Mice were head-fixed, and a customized nose cone was placed over their snout through which individual odors were delivered at a flow rate of ∼1L per minute (200A olfactometer, Aurora Scientific, Aurora, ON, Canada) with an ongoing charcoal filter vacuum system (Hydrobuilders Inc.) was used to evacuate any residual odors. Liquid sucrose solution (10% sucrose, 0.03% NaCl in water, ∼2-3 µL per presentation) was delivered via a solenoid-gated gravity feed into a spout set in front of the mouth. Contact with a lick spout positioned in front of the mouth was measured using a capacitive touch MPR121 sensor (SparkFun, Boulder, CO). Mild electric shocks were delivered via a custom-made tail cuff driven by a precision animal shocker (Coulbourn Instruments, Holliston, MA). Stimulus delivery and sensor reading were controlled by an Arduino Mega with custom circuit boards (adapted from OpenMaze.org) and recorded via CoolTerm software.

Each session consisted of 30 trials per trial type, pseudorandomly interleaved. The inter-trial interval between subsequent cues was chosen as a random sample from a uniform distribution between 8 and 17 seconds. Animals were trained on this paradigm for ∼7 consecutive days, with day 0 as a habituation day where animals only received odor presentations, day 1 labeled “Early”, and the final day of training, “Late”, where animals received all three trial types. Three neutral odors served as conditioned stimulus cues (+carvone, 2-heptanone, o-toluidine) with cue contingencies counterbalanced across mice. Presentation of the CS+ or CS-odor cue was presented for 2 seconds, followed by a 2-second trace period and subsequent delivery of either a reward = ∼2-3 µl 10% sucrose solution, tail shock = 0.1 mA 250 ms duration, or nothing (CS-trials).

### Data analysis

For statistical analyses and figures, calcium event activity was separated into 1-second and 2-second bins, and average activity during each bin was used. All statistical analyses were two-sided. Data distribution was assumed to be normal, but this was not formally tested. No statistical methods were used to pre-determine sample sizes, but our sample sizes are like those reported in previous publications^3,4^. Average calcium event rates were z-scored per trial per neuron and averaged across trials. Heatmaps and line plots were generated using calcium transients where all trials for a neuron were z-scored and baseline subtracted using data from −3 to −2 seconds prior to odor onset. To identify cells whose activity was modulated during specific epochs (for example, CS+ period and US period), for each trial the average event magnitude during the 2-second epoch was compared to a 2-second baseline period prior to cue onset for that trial and compared using a two-tailed Mann-Whitney U test, and False Discovery Rate was applied to correct for multiple comparisons. Cells with an adjusted *P* < 0.05 were classified as selective.

#### Population decoding

Analysis was done as previously described elsewhere^3,4^. A linear decoder was used to discriminate activity patterns into two discrete categories:

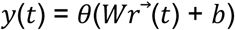

where *y* is the predicted label of the population activity pattern *r^→^* recorded at time *r^→^* and takes two values corresponding to two classes of patterns to decode (for example, two odor identities), *W* is the vector of weights assigned to each cell, and *b* is a bias term constant. Decoding parameters were attained via a supervised learning protocol with labeled data and used a support vector machine (SVM) with a linear kernel (python/scikit/linearSVC) and regularization (C) set to 1 (i.e., default value). Results are reported as the generalized performance of the decoder using cross-validation. When multiple categories were involved, e.g., more than two trial types, multiple linear decoders were trained on pairs of discrete categories combined using majority-based error correction codes. We defined the patterns of calcium activity by computing the mean event rates for each individual cell during one-second time bins. Pseudo-population recordings were generated by combining cell data across multiple animals/FOVs. For decoding, one-half of trials were randomly selected from each class and pseudopopulation activity from these trials was used to train the decoder, while the remaining held-out half was used to evaluate the decoder’s generalization performance. When comparing decoding accuracy between neural populations of different sizes, we trained our decoder on a random subsample of cells from the more numerous population equal to that of the smaller population. We repeated the operation 10 times and then combined the cross-validated decoding accuracies of all random choices together to get a single sample of decoding accuracies (i.e., a single data point reflecting the mean of 1 iteration). A two-sided Mann-Whitney U Test was used to compare decoding accuracies between groups, and Bonferroni correction was used for multiple comparisons. Decoding line plots for each data point represent the decoding accuracy for the 1-second time bin ending at that point. For example, a data point at 2 seconds after odor represents decoding accuracy using activity extracted from 1–2 seconds after odor onset.

### 1-photon imaging

Ca^2+^ videos were recorded and processed with Inscopix acquisition and preprocessing software (Inscopix, Palo Alto, CA), and triggered with a TTL pulse from EthoVision XT 10 and Noldus IO box system to allow for simultaneous acquisition of Ca^2+^ and behavioral videos for EPM and OFT. Ca^2+^ videos were acquired at 10 frames per second. An optimal LED power was selected for each mouse based on GCaMP expression in the FOV (pixel values), and the same LED settings were used for each mouse throughout the series of imaging sessions; only a single FOV was imaged per animal. Videos were spatially downsampled 2x, cropped, and band-passed filtered before Turboreg motion correction relative to a single reference frame.

Cell segmentation and calcium transient time series data were extracted using Constrained Non-negative Matrix Factorization for microendoscopic data (CNMFe), a semi-automated algorithm optimized for GRIN lens Ca^2+^ imaging to denoise, deconvolve, and demix calcium imaging data (Zhou et al., 2018). Putative neurons were manually inspected for appropriate spatial properties and Ca^2+^ dynamics and were visually checked against the corresponding motion-corrected video in ImageJ. Ca^2+^ transient events were extracted using the OASIS algorithm (Friedrich et al., 2017) embedded within CNMFe. We used calcium transients for all analyses unless otherwise noted. Denoising (CNMFe) and deconvolution (OASIS) steps were applied. Finally, the extracted calcium transient time series data were analyzed against DLC tracking data using custom Python scripts.

### Data analysis

Extracted Ca^2+^ and behavioral timestamps resolved by DLC were then analyzed using custom Python scripts. Heatmaps were created by averaging z-scored activity over the entire recording session for each neuron. To calculate population-average activity, raw Ca2+ values for each neuron were averaged across all time bins in which the animal was in either the open or closed arms of the EPM. Transition entry plots were calculated by taking each neuron’s activity baseline z-scored from −5 to −3 seconds prior to arm entry, then averaged. AUC values were calculated by taking the mean activity of each neuron and integrating it over 5 seconds after animals transitioned into the open arms. Similar was done for Head Dip plots using a −5 to −4 second baseline to z-score. For visualization only, transients were then convolved with an alpha kernel with a tau of 0.75 over 2 second window. EPM arm selectivity was determined first by temporally shuffling Ca^2+^ data for each individual neuron 1000 times to generate a null distribution. Then each neuron’s activity in each zone was subtracted from each other for the actual data and shuffled. A cell was considered selective for a zone if the calcium activity difference between the zones exceeded a 3 SD threshold in the shuffled distribution. Group comparisons were made using a χ^2^ test. For decoding analysis, first a pseudopopulation of neurons was randomly subsampled from each group, and balanced behavioral epochs were randomly subsampled with a minimum of 3 seconds bouts in each zone. The neural activity during the first 2 seconds after entry into either the closed or open-center of the maze was then used to train a linear SVM decoder (50:50 train-test split cross-validated with forward and backward accuracies) and shuffled data (shuffling arm labels 10 times) and repeated 10 times to generate final decoding accuracies. Accuracies were corrected for multiple comparisons using Bonferroni. Using the same pseudopopulation data, Principal Component Analysis was conducted using scikit-learn packages for linear dimensionality reduction fitted to a model with 5 components. Mahalanobis distance was calculated by computing the inverse covariant matrix using the first 3 PCA components and zones were subtracted from each other bootstrapped 1000 times to generate a distribution and averaged. Finally, silhouette scores were generated by calculating the intracluster mean distance and the nearest-cluster mean distance for each time bin in each zone to obtain the silhouette coefficients. Silhouette coefficients were then averaged across each time bin from each zone and compared between groups.

### Chronic Neuropixels Recordings

#### Implantation

Implants consisted of Neuropixels 2.0 4-shank probes that were assembled in a modified version of a previously published model (“Apollo implant”, Bimbard et al. 2024) with custom 3D printed parts. Probes were soldered with a silver ground wire then fixed to payload holder with Lock-ite before placing in the rest of the implant and secured with M1 screws (McMaster-Carr). In addition to the general stereotaxic procedures described above, the skull was cleaned and scored, and a thin layer of Vetbond was applied on the surface. A craniotomy of at least 1 mm in width was made centered around AP −0.420 mm, and ML 0 mm, and another hole was made above cerebellum for ground wire insertion, and finally a skull screw was placed for stabilization. Crainotomy was cleaned several times with sterile saline before insertion of probe. Probe shanks were then dipped in DiI (1,1’-Dioctadecyl-3,3,3’,3’-Tetramethylindocarbocyanine Perchlorate, Thermofisher) at least 5 times to ensure adequate coverage, then the probe was slowly lowered onto the brain tissue surface before applying Duragel. Finally, probe was lowered to a final DV −4.20 mm from bregma, before securing the implant with Lock-tite and metabond. Recovery of probe was achieved after recordings under anesthesia for reuse. Animals were then transcardially perfused with 4% PFA, and coronal sections were taken for imaging. Histology images were then used to map probe insertion sites using Herbs software aligned to the Allen Mouse Brain Atlas.

#### Recordings

Neural recordings were recorded with PXIe acquisition via SpikeGLX software at 30 kHz and data were filtered from 0.1 Hz to 5 kHz and common averaging referencing across channels. Behavior videos were recorded with a high-speed camera (Alvium 1800 U-158, Allied Vision) at 50 Hz with a 167 µs exposure and synchronized to neural recordings with external triggers. An infrared light was placed above the EPM. Tracking was achieved using DeepLabCut, followed by custom Python scripts.

### Data analysis

Neuropixels action potential signals were preprocessed with SpikeInterface and spike-sorted offline using Kilosort 4. Clusters were filtered for at least 1000 spikes over the recording session, then manually validated using Phy. Only well-isolated clusters (putative single units that are classified as ‘Good’ using Phy) were analyzed. Finally, data was curated using custom Python scripts with standard packages. To calculate arm selectivity, each neurons firing rates were binned in 20 ms to match camera frame rate for precise behavioral entrance into an arm, then z-scored over the entire recording. Arm selectivity was determined first by temporally shuffling each unit’s z-scored firing rate data for each individual neuron 1000 times to generate a null distribution. Then each neuron’s activity in each zone was subtracted from the others for both the real and shuffled data. A cell was considered selective for a zone if the activity difference between the zones exceeded a 3 SD threshold in the shuffled distribution. Spatial heatmaps were generated by binning EPM into 2.5 cm blocks; then, each unit’s spike times were summed and divided by the time spent in each bin to get firing rates. Heatmaps for open selective neurons were generated by binning unit firing rates into 250 ms blocks, then baseline z-scored −3 to −1 seconds prior to open arm entry. Head dip selective neurons were identified similarly to open arms, except head dip firing rates were subtracted from an equal number of random non-head dip bouts, then compared to differences from shuffled time bins from across the session 1000 times to generate the null distribution. Any neuron that had a +/- 3 SD threshold activity difference compared to the null distribution was considered selective. Head Dip heatmaps were generated using 200 ms binned firing rates, then baseline z-scored to −1s to −0.1s prior to the start of head dip. Finally, for the decoding zone from neural activity, behavioral epochs subsampled to balance both the open and closed arms across animals, trained a linear SVM using stratified k-fold with five folds, and compared those accuracies to a shuffled (generated by shuffling arm label 1000 times) and repeated 10 times to get average accuracies. Statistics were determined with a Mann-Whitney U two-sided test.

## Notes

### Competing Interest Statement

The authors have declared no competing interest.

